# A singleton NLR of recent origin causes hybrid necrosis in *Arabidopsis thaliana*

**DOI:** 10.1101/2020.05.18.101451

**Authors:** A. Cristina Barragan, Maximilian Collenberg, Jinge Wang, Rachelle R.Q. Lee, Wei Yuan Cher, Fernando A. Rabanal, Haim Ashkenazy, Detlef Weigel, Eunyoung Chae

## Abstract

Hybrid necrosis in plants arises from conflict between divergent alleles of immunity genes contributed by different parents, resulting in autoimmunity. We investigate a severe hybrid necrosis case in *Arabidopsis thaliana*, where the hybrid does not develop past the cotyledon stage and dies three weeks after sowing. Massive transcriptional changes take place in the hybrid, including the upregulation of most NLR disease resistance genes. This is due to an incompatible interaction between the singleton TIR-NLR gene *DANGEROUS MIX 10* (*DM10)*, which was recently relocated from a larger NLR cluster, and an unlinked locus, *DANGEROUS MIX 11 (DM11)*. There are multiple *DM10* allelic variants in the global *A. thaliana* population, several of which have premature stop codons. One of these, which has a truncated LRR domain, corresponds to the *DM10* risk allele. The *DM10* locus and the adjacent genomic region in the risk allele carriers are highly differentiated from those in the non-risk carriers in the global *A. thaliana* population, suggesting that this allele became geographically widespread only relatively recently. The *DM11* risk allele is much rarer and found only in two accessions from southwestern Spain – a region from which the *DM10* risk haplotype is absent – indicating that the ranges of *DM10* and *DM11* risk alleles may be non-overlapping.

## Introduction

Hybrid necrosis, a predominant form of hybrid incompatibility, is caused by conflicting elements of the plant immune system originating from different parental accessions. These pairwise deleterious epistatic interactions usually involve at least one nucleotide binding site-leucine-rich repeat (NLR) protein (Bomblies et al. 2007; Alcázar et al. 2009; Yamamoto et al. 2010; Chae et al. 2014; Sicard et al. 2015; Deng et al. 2019). NLRs function as intracellular immune receptors homologous to NOD/CARD genes in animals, and play a major role in plant innate immunity (Maekawa et al. 2011; Jones et al. 2016). The constant co-evolutionary arms-race between plants and their pathogens has led to a high diversification of many elements of the plant immune system, including NLRs (Jones and Dangl 2006; Dodds and Rathjen 2010). Hybrid necrosis can be viewed as collateral damage resulting from excessive sequence diversity, a phenomenon that may limit the possible NLR allele combinations found in an individual plant (Chae et al. 2014).

Plant NLRs are multidomain proteins usually composed of N-terminal Toll/interleukin-1 receptor (TIR), coiled-coil (CC) or RESISTANCE TO POWDERY MILDEW 8 (RPW8) domains, a central nucleotide-binding site (NBS) and C-terminal leucine-rich repeats (LRRs) (Meyers et al. 2003; Shao et al. 2016). The N-terminal domain is usually thought to be involved in signal transduction, while the NBS domain can act as a molecular ON/OFF switch (Bentham et al. 2017). The LRR domain is highly variable and consists of multiple repeats of 20-30 amino acid stretches that are often responsible for direct or indirect pathogen effector recognition as well as NLR auto-inhibition (Ade et al. 2007; Krasileva et al. 2010; Steinbrenner et al. 2015).

Approximately half of all NLRs in a given *A. thaliana* accession are found in multi-gene clusters, which are unevenly distributed across the genome (Meyers et al. 2003; Van de Weyer et al. 2019). Tandem duplication events are common in NLR clusters, and duplicate genes are a major source of genetic variation, since they often experience relaxed selection and enable neofunctionalization (Ohno 1970; Force et al. 1999; Lynch and Conery 2000; Conant and Wolfe 2008). Sequence homogenization through intergenic exchange among cluster members is avoided when an NLR gene is translocated away from its original cluster to an unlinked genomic region, thereby preserving its original function or potentially developing a new one (Baumgarten et al. 2003; Leister 2004). For NLRs, neofunctionalization of duplicated or translocated genes can expand the repertoire of pathogen effectors an individual plant is able to recognize (Botella et al. 1998; Michelmore and Meyers 1998; Holub 2001; Kim et al. 2017).

Genome-wide analysis of structural variation across eight high-quality *A. thaliana* genomes identified rearrangement hot spots coinciding with numerous multi-gene NLR clusters (Jiao and Schneeberger 2020), including the previously described *DANGEROUS MIX* (*DM*) loci, which are causal for hybrid necrosis (Bomblies et al. 2007; Chae et al. 2014). This raises the possibility that accelerated evolution associated with genomic rearrangements contribute to the generation of incompatibility alleles, pointing to genomic architecture as a driver of hybrid incompatibility. So far, over a dozen NLR loci with hybrid necrosis alleles are known from multiple plant species, yet none of them is a singleton NLR, even though singletons account for about a quarter of NLRs in different species (Jacob et al. 2013). Most, but not all, well-characterized singleton NLRs, such as *RPM1* and *RPS2* in *A. thaliana*, show ancient balanced polymorphisms that maintain active and inactive alleles at intermediate frequencies in natural metapopulations (Caicedo et al. 1999; Stahl et al. 1999; Mauricio et al. 2003; Allen et al. 2004; MacQueen et al. 2016). Thus, with less functional diversity, and beneficial alleles often being relatively common, one would indeed expect that singleton NLRs are underrepresented among hybrid necrosis loci.

Here, we are investigating a case of severe hybrid necrosis, where hybrid plants do not develop past the cotyledon stage, become necrotic, and die three weeks after sowing. Extensive transcriptional changes occur in the hybrid, including the induction of most NLR genes. Through a combination of QTL analysis and GWAS, we identified two new incompatibility loci, *DANGEROUS MIX 10* (*DM10*), a TIR-NLR on chromosome 5, and *DM11*, an unlinked locus on chromosome 1, as causal for incompatibility. *DM10* is an unusual hybrid incompatibility locus because it is a singleton NLR that arose after *A. thaliana* speciation through interchromosomal transposition from the *RLM1* cluster, which confers resistance to blackleg disease in multiple *Brassica* species and *A. thaliana* (Richly et al. 2002; Staal et al. 2006; Guo et al. 2011). The causal allele has a premature stop codon that removes the C-terminal quarter of the protein, highlighting that substantial truncations of the coding region do not necessarily indicate loss of function.

## Results

### A particularly severe case of hybrid necrosis: Cdm-0 x TueScha-9

Eighty *A. thaliana* accessions have previously been intercrossed with the goal of identifying hybrid incompatibility hot spots (Chae et al. 2014). A particularly severe case was observed in the crosses between Cdm-0 and five other accessions: TueScha-9, Yeg-1, Bak-2, ICE21 and Leo-1. The F_1_ progeny of these two parents did not develop past the cotyledon stage, even at temperatures that suppress hybrid necrosis in most other cases (Chae et al. 2014), and severe necrosis developed during the second week after sowing, followed by complete withering in the third week (**Fig 1A**).

**Fig 1.**
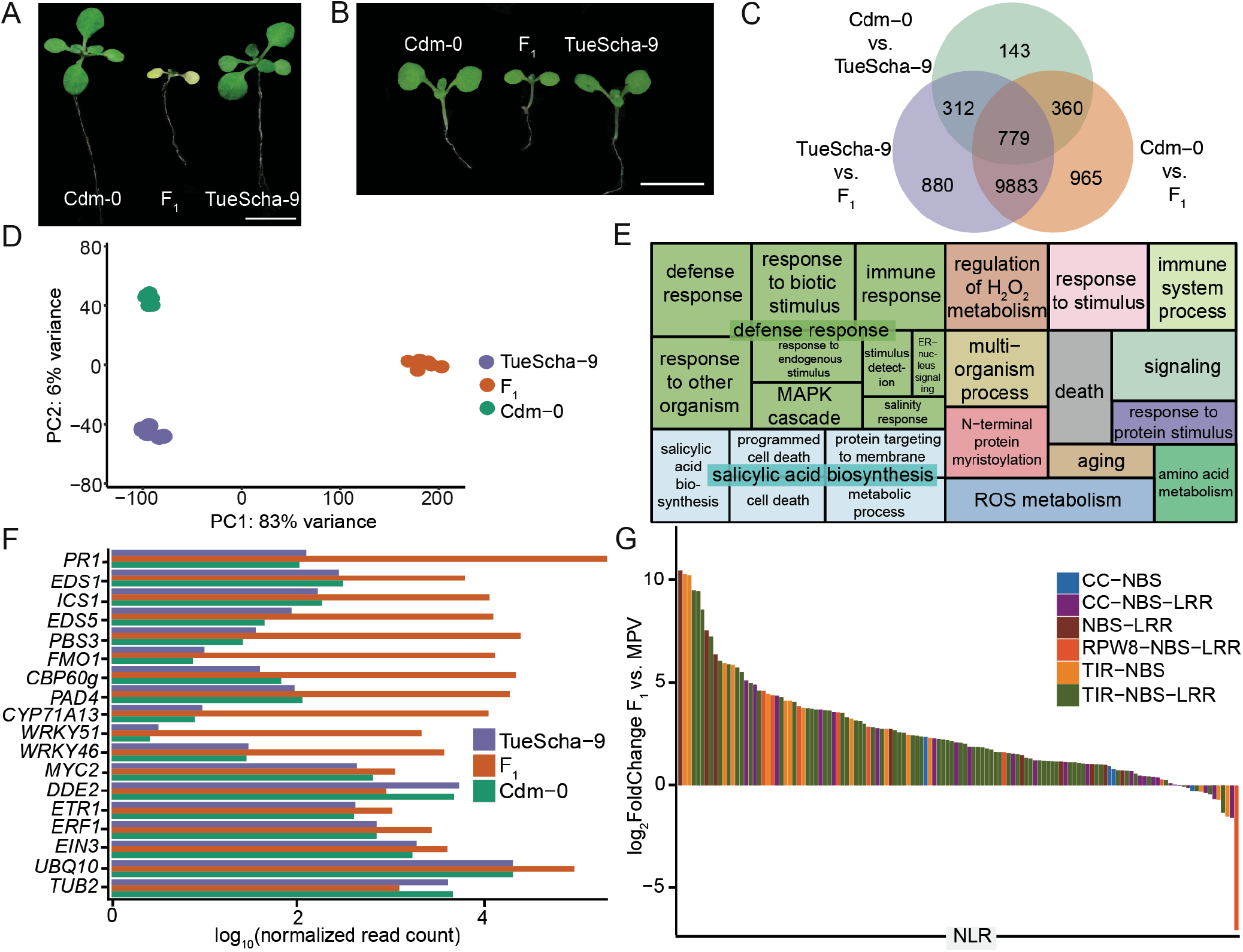
RNA-seq analysis of Cdm-0 x TueScha-9 hybrids. **A**. At 21 days, the Cdm-0 x TueScha-9 F_1_ hybrid is necrotic. Plants were grown at 16°C. Scale bar represents 1 cm. **B.** Examples of a 10-day old Cdm-0 x TueScha-9 F_1_ hybrid and parental accessions harvested for RNA-seq. Plants were grown at 23°C. Scale bar represents 1 cm. **C.** Intersection of DEGs between the F_1_ hybrid and parents. **D.** PCA of gene expression values. The main variance is between the F_1_ hybrid and parents. Each dot indicates one biological replicate, with six per genotype. **E.** REVIGO Gene Ontology treemap. Size of the square represents −log_10_(*p* value) of each GO term. **F.** −log_10_(normalized read count) of defense-related marker genes of the hybrid and the parents. **G.** NLR expression changes between the F_1_ hybrid and the MPV, with 128 significantly (|log_2_ FoldChange| >1, padj value < 0.01) differentially expressed in at least one genotype comparison.

To obtain insights into the transcriptional changes in the hybrid, we performed RNA-seq on the parental accessions Cdm-0 and TueScha-9, as well as in F_1_ hybrid plants 10 days after germination, when the hybrid was already slightly stunted, but before there were visible signs of necrosis (**Fig 1B**). We observed massive transcriptional changes, in which around half of all 20,000 detectable genes (**Fig S1A**) were differentially expressed in the hybrid when compared to either parent (**Fig 1C**, **S1B**, **Table S1**). This represents one third of the entire *A. thaliana* transcriptome (Klepikova et al. 2016). A principal component analysis (PCA) showed that most of the variance in gene expression is driven by the difference between the parents and the hybrid (PC1: 83 %) (**Fig 1D**). In addition, we generated in silico hybrids (see Methods) and compared these with the biological F_1_ hybrids through a PCA. This confirmed that gene expression in the F_1_ hybrid is not an additive result of expression in the two parental accessions (**Fig S1C**). Next, we carried out a Gene Ontology (GO) analysis using the top 1,000 differentially expressed genes (DEGs) between the F_1_ hybrid and the MPV (**Table S2**). “Defense response” and “salicylic acid biosynthesis” were the categories with the highest number of DEGs in the hybrid versus MPV comparison (**Fig 1E**, **Table S2**).

Since the F_1_ hybrid displayed signs of an increased pathogen defense response, we analyzed the expression of a set of marker genes for defense-associated phytohormones such as jasmonic acid (JA), salicylic acid (SA) and ethylene (ET) (Papadopoulou et al. 2018), as well as early pathogen response genes induced by both cell surface receptors and NLRs (Ding et al. 2020) (**Table S3**). Genes involved in SA biosynthesis and signaling, such as *EDS1, ICS1*, *EDS5*, *PAD4*, *PBS3, CBP60* and *FMO1*, were strongly overexpressed in F_1_ hybrid plants, in concordance with the GO analysis, as was the SA-induced camalexin biosynthesis gene *CYP71A13*. The expression of genes encoding transcription factors *WRKY46* and *WRKY51* and of the late immune response gene *PR1* was also increased in the hybrid (**Fig 1F**, **Table S3**). In contrast, the expression of genes required for JA-mediated resistance, such as *MYC2* or *DDE2*, or genes involved in ET signaling, such as *ETR1, ERF1* and *EIN3*, changed to a lesser extent in the F_1_ hybrid, similar to control genes *UBQ10* and *TUB2* (**Fig 1F**, **Table S3**).

Since an increase in NLR expression has been linked to autoimmunity (Stokes et al. 2002; Mackey et al. 2003; Palma et al. 2010; Lai and Eulgem 2018), and since some NLRs are upregulated when SA levels rise (Shirano et al. 2002; Yang and Hua 2004; Tan et al. 2007; Mohr et al. 2010; MacQueen and Bergelson 2016), we set out to investigate NLR expression levels in the hybrid. Out of a set of 166 NLRs found in the Col-0 genome, 150 were expressed in at least one of the three genotypes studied, and 128 were significantly (|log_2_FoldChange| >1, padj value < 0.01) differentially expressed in at least one genotype comparison (**Fig S2D**, **Table S4**). From these 128 NLRs, all but one were differentially expressed when comparing the hybrid with either parental accession (**Fig S1D**). NLRs were mostly upregulated in the F_1_ hybrid: of the 95 NLRs with significant expression changes in the hybrid versus the MPV, all but three were overexpressed (**Fig 1G**, **S1E**, **Table S1**). When the F_1_ hybrid was compared to the parents, the expression of individual NLRs largely followed the same pattern (**Fig S1E**-**G**), this was not the case when comparing the two parents (**Fig S1H**). The fraction of genes overexpressed in the F_1_ hybrid was similar for the different NLR classes as well as between singleton and clustered NLRs (**Fig 1G**, **S1F-H**, **Table S3**).

### QTL mapping of *DM10* and *DM11* in a triple-hybrid cross

Having found that a very large fraction of NLR genes is upregulated in the Cdm-0 x TueScha-9 hybrid, we wondered whether hybrid necrosis in this case was due to global NLR regulators (Li et al. 2009; Zhai et al. 2011; Shivaprasad et al. 2012; Gloggnitzer et al. 2014; Sicard et al. 2015), or to NLRs, as in other hybrid necrosis cases. We therefore proceeded to map the underlying causal loci via quantitative trait locus (QTL) analysis. Since the F_1_ hybrid seedlings died very young, we could not directly generate a segregating F_2_ mapping population (Bomblies et al. 2007; Chae et al. 2014; Barragan et al. 2019). Instead, we designed a triple-hybrid cross (Cooper et al. 2019) and first generated two sets of heterozygous plants by crossing Cdm-0 and TueScha-9 separately to a third, innocuous background, the Col-0 reference accession. We then intercrossed these Cdm-0/Col-0 and TueScha-9/Col-0 plants (**Fig 2A**). In the resulting pseudo-F_2_ generation, we collected both normal and necrotic plants and individually genotyped them by RAD-seq (Rowan et al. 2017). For QTL mapping, we focused on polymorphic markers between Cdm-0 and TueScha-9, including markers overlapping with the Col-0 reference (**Fig 2B**), and also analyzed polymorphic markers for each accession independently (**Fig S2A**, **B**). We identified two genomic regions that interact epistatically to cause the severe hybrid necrosis phenotype. We called the QTL on chromosome 5 (23.35 to 24.45 Mb) *DM10*, and the QTL on chromosome 1 (21.55 to 22.18 Mb) *DM11*. Both intervals contained NLRs but no clear candidates for global NLR regulators, so we chose to focus on NLR genes. In the *DM10* mapping interval, one NLR was present, At5g58120, while the *DM11* interval was NLR-rich and encompassed 10 NLRs in Col-0 (**Table S5**). Loci in the interval included the highly polymorphic *RPP7* cluster of CC-NLR genes (McDowell et al. 2000; Guo et al. 2011; Li et al. 2020), as well as the two CC-NLR singleton genes, *CW9* (At1g59620) and At1g59780 (Meyers et al. 2003). To identify potential differences between Col-0 and Cdm-0 in the *DM11* interval, we generated a PacBio long-read-based genome assembly of this accession (**Table S6**). Notably, most chromosome arms were assembled in single contigs, including the long arm of chromosome 1, where the *DM11* mapping interval is located (**Fig S3**). Since at the time the full Cdm-0 annotation was not yet available, we manually annotated homologs of NLR genes corresponding to the genomic region that spans from At1g56510 to At1g64070 in Col-0, which includes the *DM11* mapping interval as well as neighboring NLRs. Like Col-0, Cdm-0 carries groups of both clustered and singleton NLRs, adding up to a total of 21 NLRs, compared to 28 NLRs in Col-0 (**Fig 2C**, **Table S5**).

**Fig 2.**
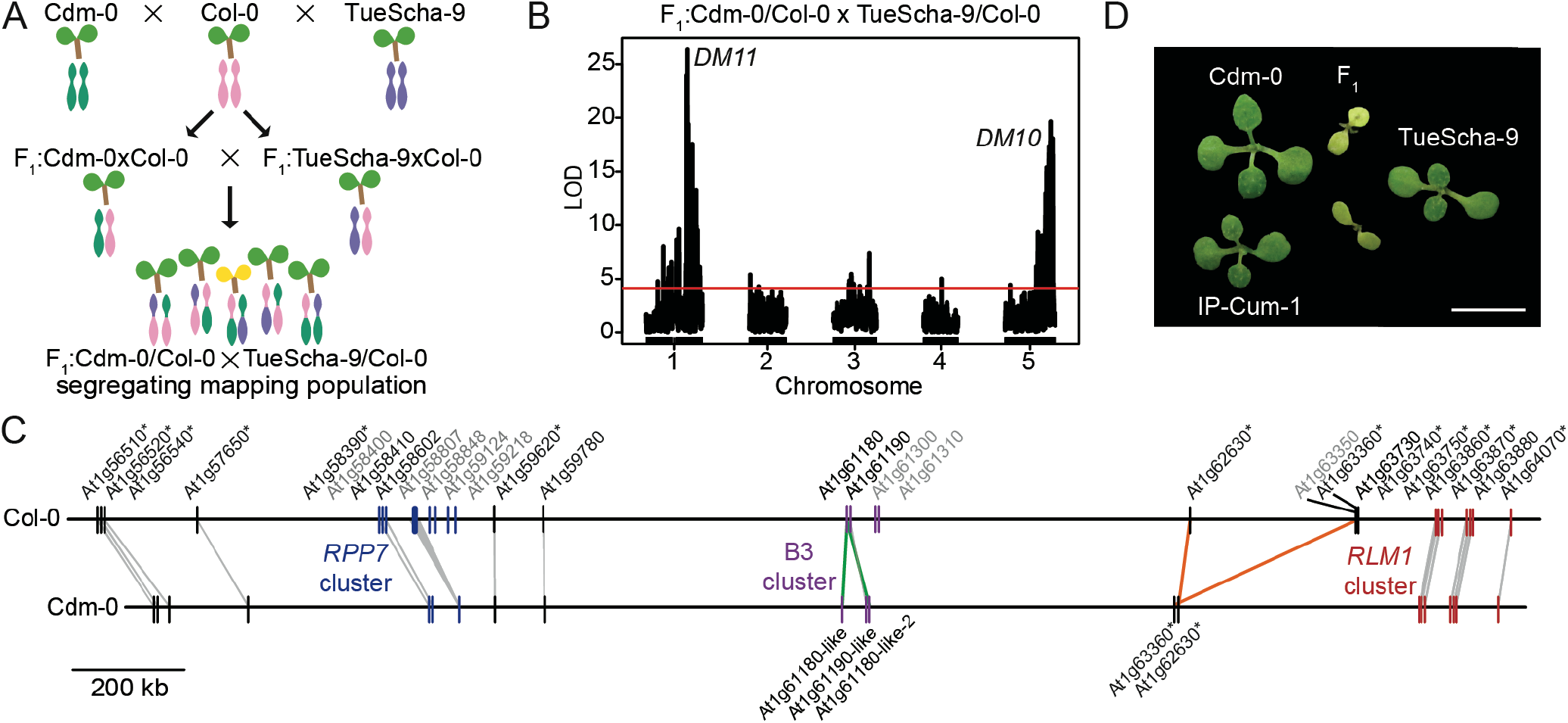
QTL mapping with a triple-hybrid cross. **A.** Creation of a Cdm-0 x TueScha-9 mapping population. **B.** QTL analysis from polymorphic Cdm-0 and TueScha-9 markers. QTL peaks are found on chromosome 5 (23.35-24.45 Mb), *DM10*, and chromosome 1 (21.98-22.04 Mb), *DM11*. The horizontal lines indicate 0.05 significance threshold established with 1,000 permutations. **C.** Comparison and distribution of candidate *DM11* NLR genes between At1g56510 and At1g64070 on chromosome 1. Gene IDs in grey are present in Col-0 but not in Cdm-0, gene duplications are marked in green and inversion events in orange. Asterisks indicate significant (|log_2_ FoldChange|>1, padj value < 0.01) gene expression changes in the F_1_ hybrid when compared to the MPV. **D.** Cdm-0 x TueScha-9 and IP-Cum-1 x TueScha-9 hybrids. Plants are two weeks old and were grown at 16°C. Scale bar represents 1 cm.

To pinpoint *DM11* candidate genes, we sought to identify additional accessions that had similar alleles as Cdm-0 at *DM11* candidate loci by creating Neighbor-Joining (NJ) trees (**Fig S2C**) and PCA plots (**Fig S2D**, **Table S7**), using sequences from the 1001 Genomes Project (1001 Genomes Consortium 2016). IP-Cum-1 was the accession most similar to Cdm-0 for the whole *DM11* mapping interval, and when we crossed it to TueScha-9, Cdm-0 x TueScha-9-like hybrid necrosis was observed (**Fig 2D**). Eleven other accessions that were less closely related to Cdm-0 in this genomic interval did not produce necrotic F_1_ hybrids (**Table S7**). Because accessions Istisu-1 and ICE134, like Cdm-0, lack a transposable element that is present in most *RPP7* (At1g68602) alleles (Tsuchiya and Eulgem 2013), we also crossed these two accessions to TueScha-9, but no hybrid necrosis was observed (**Table S7**). Artificial miRNAs (amiRNAs) (Schwab et al. 2006) targeting different members of the *RPP7* cluster were previously designed to perform rescue experiments (Chae et al. 2014; Barragan et al. 2019); although predicted to target all members of the Cdm-0 *RPP7* cluster, neither these nor amiRNAs targeting *CW9*^Cdm-0^ or At1g59780^Cdm-0^ suppressed hybrid necrosis (**Table S8**). Lastly, a genomic *CW9*^Cdm-0^ fragment was unable to induce hybrid necrosis when introduced into TueScha-9 (**Table S15**).

Being aware that the precision of QTL mapping in NLR-rich regions can be affected by structural variation, we also tested NLRs adjacent to the *DM11* mapping interval. The *RLM1* cluster is highly similar among Cdm-0 and IP-Cum-1, both of which carry the causal *DM11* allele in addition, some cluster members show an increased expression in the F_1_ hybrid, which is sometimes the case for causal NLRs (Bomblies et al. 2007) (**Fig S2E**). We therefore tested six of the seven *RLM1* cluster members via *Nicotiana benthamiana* co-expression with *DM10*^TueScha-9^ (see Fig 4 for cloning of causal *DM10* allele), but none induced a hypersensitive response (HR) (**Table S5**). Six accessions with a similar *RLM1* locus to that of Cdm-0 and IP-Cum-1 were crossed with TueScha-9, but no necrosis was observed (**Table S7**). Finally, because At1g57650 was strongly upregulated among *DM11* NLR candidate genes, we tested it with *DM10*^TueScha-9^ in *N. benthamiana*, but again no HR was observed (**Fig S2E**, **Table S4**, **S5**). This may indicate that *DM11* is either an NLR that was not tested, or not an NLR at all. Note that some Col-0 NLRs that had no homologs in the interval from At1g56510 to At1g64070 in Cdm-0 attracted nonspecific RNA-seq reads, most likely because there are homologs elsewhere in the Cdm-0 genome (**Table S5**).

**Fig 4.**
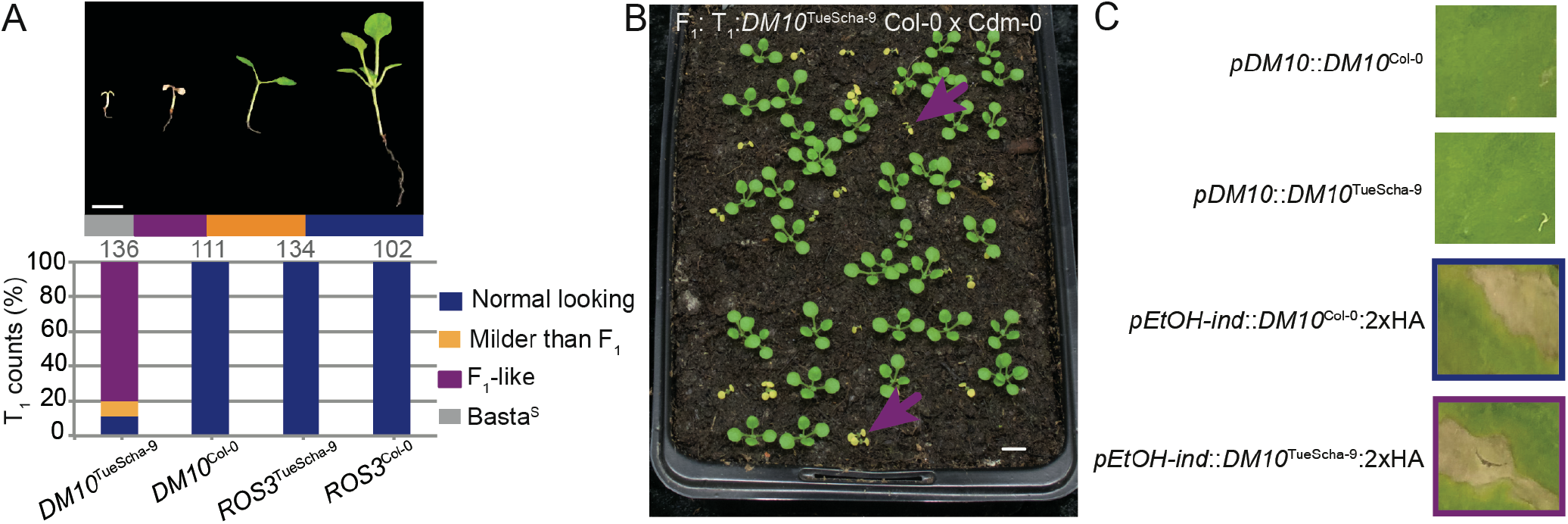
Experimental identification of *DM10*. **A.** Recapitulation of hybrid necrosis in 25-day old Cdm-0 T_1_ plants transformed with the indicated genomic fragment from TueScha-9 or Col-0. Representative phenotype and total number of T_1_ plants examined given on top. Plants were grown at 16°C. Scale bar represents 1 cm. **B.** The same *DM10*^TueScha-9^ genomic fragment as in A was introduced into Col-0, and T_1_ plants were crossed to Cdm-0. The F_1_ hybrid phenotype was recapitulated (magenta arrows). Plants were 18 days old and grown at 16°C. Scale bar represents 1 cm. **C.** Infiltration of *N. benthamiana* leaves with the indicated constructs. Overexpression of either *DM10*^TueScha-9^ or *DM10*^Col-0^ under an EtOH inducible promoter (*pEtOH-ind*) triggered cell death, while expression from their native promoter (*pDM10*) did not.

### Fine mapping of *DM10* using Genome-Wide Association Studies

In the original collection of 6,409 crosses among 80 accessions (Chae et al. 2014), four accessions in addition to TueScha-9 produced severe hybrid necrosis when crossed to Cdm-0: Yeg-1, Bak-2, ICE21 and Leo-1. Together with TueScha-9, these represent much of the Eurasian range of the species, both geographically and genetically; six of the nine previously identified admixture groups (1001 Genomes Consortium 2016) are present in these five risk accessions (**Fig 3A**, **Table S9**). Given the diversity of the five incompatible accessions, and knowing that most, but not all, causal genes for hybrid incompatibility are NLRs, we attempted to narrow down causal *DM10* candidate genes by GWAS, with Cdm-0-dependent F_1_ necrosis as a binary trait (Grimm et al. 2016). We discovered a remarkably high association between this phenotype and several closely linked markers on the bottom of chromosome 5, with corrected *p* values as low as 10^−38^. In addition, 79 SNPs showed a one-to-one association with the necrotic phenotype, resulting in −log_10_ *p* values of 0 (**Fig 3B**, **Table S10**). The markers with the strongest associations tagged three loci: At5g58120, encoding a TIR-NLR without known function*, ROS3* (At5g58130), encoding an enzyme involved in DNA demethylation (Zheng et al. 2008), and *PHOT2* (At5g58140), encoding a blue light receptor that mediates phototropism (Harada et al. 2003) (**Fig 3C**, **Table S10**). These three loci are genetically similar among the five risk accessions, yet differentiated from the other 75 accessions used for GWAS (**Fig S4A**-**C**). Looking at linkage among loci in this genomic region, we could see that, when taking all 80 accessions into account, six loci (At5g58090-Atg58140) belong to one large linkage block, in which *ROS3* and *PHOT2* are under tight linkage and the TIR-NLR At5g58120 constitutes a separate linkage block (**Fig 3D**). Notably, in the five accessions causing hybrid incompatibility, stronger linkage is observed in this region than that seen among the same markers from all 80 accessions (**Fig 3E, F**). In the risk accessions, At5g58120, *ROS3* and the proximal part of *PHOT2* form one linkage block, while SNPs located in the distal half of *PHOT2* are found in a separate linkage block, rendering At5g58120 and *ROS3* as primary candidates for causality in hybrid necrosis (**Fig 3F**).

**Fig 3.**
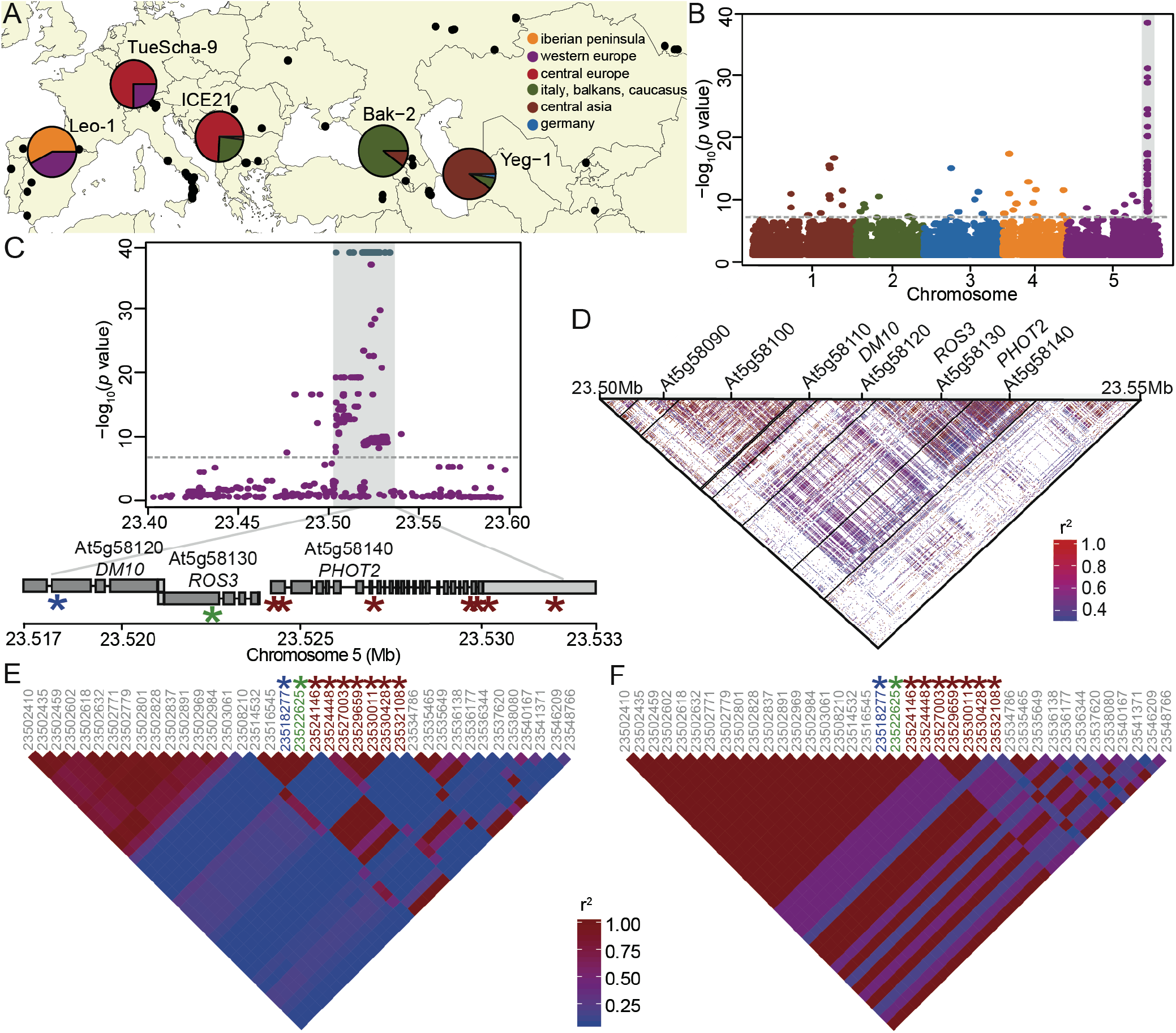
GWAS of hybrid necrosis in 80 accessions. **A.** Map of 80 accessions (black dots), with the five risk accessions colored according to 1001 Genomes admixture groups (1001 Genomes Consortium 2016). **B.** Manhattan plot for association of necrosis in Cdm-0 hybrid progeny when selfed and crossed to 79 other accessions. The significance threshold (Bonferroni correction, 5% familywise error) is indicated as a horizontal dotted line (same in C). **C.** Close-up of the region highly associated with hybrid necrosis; SNPs with a 1:1 association marked in teal. Asterisks indicate such 1:1 associations in At5g58120, *ROS3* and *PHOT2*; see also E and F. SNP positions are given in Table S10. **D.** Linkage disequilibrium (LD) across a 50 kb region in chromosome 5. Strong linkage is observed from At5g58090 to At5g58140. **E.** LD across the same 50 kb region as in D, with a subset of markers from 80 accessions crossed to Cdm-0. Asterisks indicate markers highlighted in C. **F.** LD across a 50 kb region with the same markers as in E, but for the five risk accessions only. Higher LD is seen here than in E.

### *DM10*, a singleton TIR-NLR, as cause of severe hybrid necrosis

Having candidate genes for *DM10*, we next sought to experimentally test their causality for severe hybrid necrosis. Genomic fragments of the TIR-NLR At5g58120 and *ROS3*, from both Col-0 and TueScha-9, were introduced into Cdm-0 plants. A 4.8 kb genomic fragment containing At5g58120^TueScha-9^ recapitulated the Cdm-0 x TueScha-9 hybrid necrosis phenotype (**Fig 4A**, **Table S11**). At5g58120 is henceforth called *DM10*. When *DM10*TueScha-9 was introduced into a Col-0 background and the resulting T_1_ plants were subsequently crossed to Cdm-0, we also observed the hybrid incompatibility phenotype in the F_1_ progeny (**Fig 4B**). *DM10*Col-0*, ROS3*^TueScha-9^ and *ROS3*^Col-0^ did not produce any necrosis when introduced into a Cdm-0 background (**Table S11**). We also observed that, when infiltrated in *N. benthamiana* leaves and overexpressed under an EtOH-inducible promoter, both *DM10*^Col-0^ and *DM10*^TueScha-9^ were able to trigger cell death, which was not the case when *DM10*^Col-0^ and *DM10*^TueScha-9^ were expressed under the control of their native promoters (**Fig 4C**). The cell death triggering activities under forced overexpression in the heterologous *N. benthamiana* system indicate that these NLRs are functional and competent in immune signaling.

### Prevalence and genetic differentiation of the *DM10* risk allele in the global *A. thaliana* population

To study natural variation across different *DM10* alleles, 73 alleles belonging to accessions originating from across *A. thaliana’*s native range were extracted from preliminary short- and long-read genome assemblies available in-house (**Fig S5A**, **Table S12**). A Maximum-Likelihood (ML) tree showed that there are multiple well-supported DM10 clades (**Fig S5B**), and that variation between DM10 proteins was most prevalent at the C terminal end (**Fig 5A**, **Fig S5C**, **Table S13**). Ten alleles were predicted to produce proteins truncated at three different points. Four accessions, including TueScha-9, the original *DM10* risk allele carrier, share the same stop codon (**Fig 5B**, **S5B**), removing three LRRs. Five accessions had shorter, 335 amino acid long DM10 proteins; in these, the NBS domain was truncated, lacking motifs which are important regulators of NLR activity (Bendahmane et al. 2002; Sueldo et al. 2015; Bentham et al. 2017) (**Fig 5B**, **Fig S5C**). These five accessions carrying short *DM10* alleles included Cdm-0 and IP-Cum-1, which also carry *DM11* risk alleles. This implies that the short Cdm-0-like *DM10* variants do not interact with *DM11* to produce hybrid necrosis. The shortest predicted DM10 protein, found in the Sha accession, was only 90 amino acids long and became truncated midway through the TIR-2 motif (**Fig 5B**, **Fig S5C**). The full-length DM10^Col-0^ and the truncated DM10^TueScha-9^ proteins differ only at 3% of shared sites, which is low for within-species variation among NLR alleles (Van de Weyer et al. 2019). As is typical for NLRs (Mondragón-Palomino et al. 2002; Ruggieri et al. 2014), Ka/Ks values above 1 were found in the LRR domain when comparing DM10^Col-0^ and DM10^TueScha-9^ (**Fig 5C**). In contrast, DM10^Col-0^ and DM10^Lerik1-3^, which are both full-length DM10 proteins but from different clades, are more differentiated in TIR and NB-ARC domains, although Ka/Ks values above 1 are also restricted to the LRR domain (**Fig S5D**).

**Fig 5.**
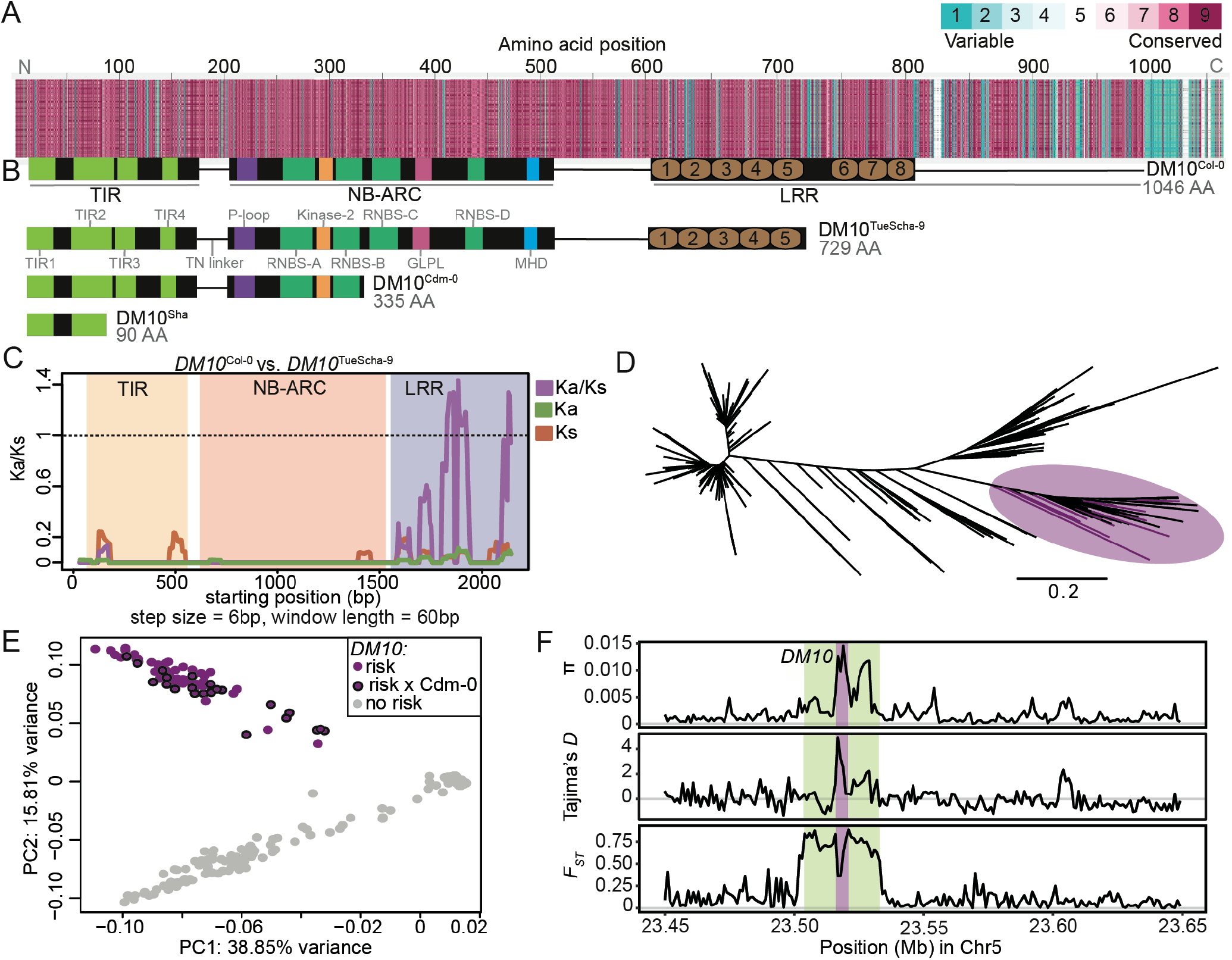
*DM10* natural variation. **A.** Amino acid alignment of 73 DM10 proteins color-coded by its conservation score (Armon et al. 2001). **B.** Comparison between different DM10 proteins (aligned with A). **C.** Ka/Ks ratios between *DM10*^Col-0^ and *DM10*^TueScha-9^. **D.** NJ tree of a region including *DM10*, *ROS3* and *PHOT2* sequences from the 1001 Genomes Project (1001 Genomes Consortium 2016). Branch lengths in nucleotide substitutions are indicated. Accessions carrying the *DM10* risk alleles group together in a branch (magenta), risk accessions crossed to Cdm-0 are highlighted. **E.** PCA. Accessions carrying the predicted *DM10* risk (magenta) versus non-risk (grey) alleles are clearly separated in PC2. Risk accessions crossed to Cdm-0 are outlined in black. **F. π**, Tajima’s *D* and *F_ST_* for *DM10* (magenta), the *DM10* linkage block comprising At5g58090-Atg58140 (green) and surrounding genomic regions.

Next, to assess how common the *DM10* risk allele is in the global *A. thaliana* population, we again turned to the 1001 Genomes collection (1001 Genomes Consortium 2016). Since *DM10*, *ROS3* and the proximal part of *PHOT2* were strongly linked in accessions carrying the *DM10* risk allele, we focused on this region, which contained 785 SNPs. In a NJ tree, all five confirmed *DM10* risk allele carriers were found in the same branch, which included 95 other accessions (**Fig 5D**, **Table S14**). In a PCA of this region, these 100 accessions were clearly separated from the rest (**Fig 5E**), which was not the case in a whole-genome PCA (**Fig S5E**), indicating that population structure is not the main driving force separating risk from non-risk allele carriers. To experimentally confirm that sequence was predictive of interaction with the *DM11* risk allele, 25 of the 100 accessions were crossed to Cdm-0 (**Fig S5B**, **Table S14**). All but IP-Alm-0 produced hybrid necrosis. Notably, while DM10 from IP-Alm-0 is 99.2% identical with DM10^TueScha-9^, it does not have the LRR truncation (**Fig S5B**). This implies that the truncation in *DM10* risk alleles is likely responsible for incompatibility, and not individual amino acid changes. Ten random accessions not predicted to carry the *DM10* risk allele were crossed to Cdm-0 as a control; as expected, none produced hybrid necrosis (**Table S14**). Similarly, we investigated how common the other two DM10 truncations are in the global *A. thaliana* population. The shortest Sha-like *DM10* allele was found in 29 accessions, while the Cdm-0-like truncation was more common, although not as common as the *DM10* risk allele, and was found in 67 accessions (**Table S14**).

In a 200 kb region around *DM10*, nucleotide diversity (**π**) was highest, up to 0.015, in the distal half of *DM10*, encoding the more polymorphic LRR domain (**Fig 5F**). However, in comparison with other TIR-NLRs present in most or all accessions, overall *DM10* nucleotide diversity was not uncommon (Van de Weyer et al. 2019). Values for Tajima’s *D* reached 4.8 in the proximal half of *DM10,* hinting at multiple *DM10* alleles being prevalent at intermediate frequencies in the global *A. thaliana* population, as is often the case for NLRs (Caicedo et al. 1999; Stahl et al. 1999; Bakker et al. 2006; Karasov et al. 2014). Lastly, the fixation index (*F_ST_*) between 98 accessions with predicted *DM10* risk alleles (excluding IP-Alm-0 and RAD-21, which did not have truncated LRR domains) and 1,037 non-risk allele carrying accessions, peaked at 0.88 across the *DM10* linkage block (**Fig 3D**-**F**, **Fig 5F**). This was the only peak detected both across the entire chromosome 5 (**Fig S5F**) and the whole genome. Inside this block, a drop in *F_ST_* is seen over the proximal half of *DM10*, which is consistent with this region being similar between risk and some non-risk alleles (**Fig 5C**).

Taken together, these results show that there are multiple *DM10* alleles in the global *A. thaliana* population, three of which are predicted to result in truncated proteins due to the presence of premature stop codons, one of which is the *DM10* risk allele. Notably, the *DM10* risk allele is not only relatively common and genetically differentiated in our GWAS population, but also in the global *A. thaliana* population.

### No documented co-occurrence of *DM10* and *DM11* risk alleles in the global *A. thaliana* population

Looking at the geographical distribution of accessions carrying different *DM10* alleles with premature stop codons, we observed that both the Cdm-0-like *DM10* allele as well as the risk *DM10* allele were found at similar densities throughout *A. thaliana*’s native range, while the Sha-like *DM10* allele was mainly found towards the eastern part of the species’ distribution (**Fig 6A**, **Table S14**). In the case of the *DM10* risk allele, the one exception to where this allele was found, was the southwestern part of Spain and Portugal, even though *A. thaliana* has been heavily sampled in this region (1001 Genomes Consortium 2016). The fact that the only two *DM11* risk carriers identified so far, Cdm-0 and IP-Cum-1, are found in southwestern Spain may indicate that the *DM10* and *DM11* risk alleles do not geographically co-occur (**Fig 6B**).

**Fig 6.**
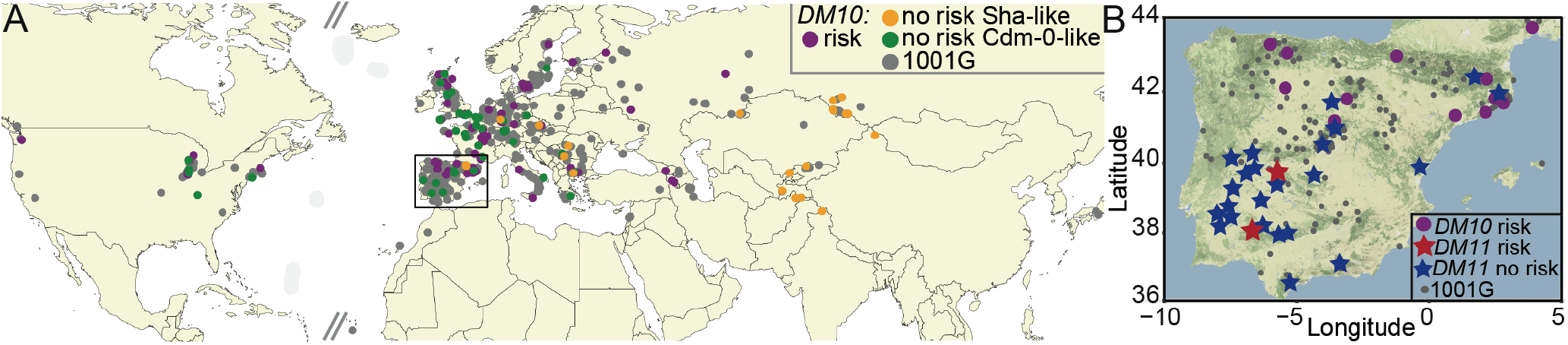
Geographical distribution of *DM10* and *DM11* alleles. **A.** Geographical distribution of 1001 Genomes Project accessions (1001 Genomes Consortium 2016) carrying the *DM10* risk (magenta), non-risk (grey) alleles, Sha-like non-risk (orange) and Cdm-0-like non-risk (green) alleles. Rectangle zooms into the region shown in B. **B.** Distribution of 1001 Genomes Project accessions (grey) in Spain and Portugal, carrying the *DM10* (magenta) and *DM11* (red) risk alleles, as well as accessions carrying *DM11* non-risk alleles (blue) which were crossed to TueScha-9.

To provide additional support for this assertion, we first attempted to identify more *DM11* risk carriers in Spain and Portugal. We crossed TueScha-9, a *DM10* risk allele carrier, to 24 accessions from these two countries, which were found at different geographical distances from the two *DM11* risk carriers Cdm-0 and IP-Cum-1, as well as from accessions carrying the *DM10* risk allele (**Table S7**). No hybrid necrosis was observed in any of the resulting F_1_ progeny (**Fig 6B**, **Table S7**). This, together with our aforementioned attempts to find additional *DM11* carriers among accessions that are closely related in the *DM11* genomic region to Cdm-0 and IP-Cum-1, indicates that the *DM11* risk allele is rare and potentially only found in southwestern Spain, a region where the *DM10* risk allele appears to be absent.

### Origin of the *DM10* NLR singleton locus through a recent interchromosomal relocation event out of the *RLM1* cluster

In the *A. thaliana* Col-0 reference genome, we identified nine NLR genes closely related to *DM10*. Seven of these make up the *RLM1* cluster on chromosome 1, and two others, At2g16870, At4g14370 are dispersed singletons. In the related species *Arabidopsis lyrata* and *Brassica rapa*, we identified a further 20 *DM10/RLM1* homologs (**Fig 7A**). As in *A. thaliana*, the *RLM1* cluster in these two species underwent within-species duplication and inversion events (**Fig 7B**). Most *RLM1* members from *A. thaliana* have a clear one-to-one homolog in *A. lyrata*, so the expansion of the *RLM1* cluster must have occurred before the two species diverged (Beilstein et al. 2010). The *A. lyrata* homologs of At2g16870 and At4g14370, 480565 and 493465, are found in a different chromosome than the main *RLM1* cluster (**Fig 7C**). This is not the case for the *DM10* homolog from *A. lyrata*, 875509, which is located inside the main *RLM1* cluster (**Fig 7C**). This indicates that *DM10* was relocated away from the main *RLM1* cluster to another chromosome and that this occurred after *A. lyrata* and *A. thaliana* diverged.

**Fig 7.**
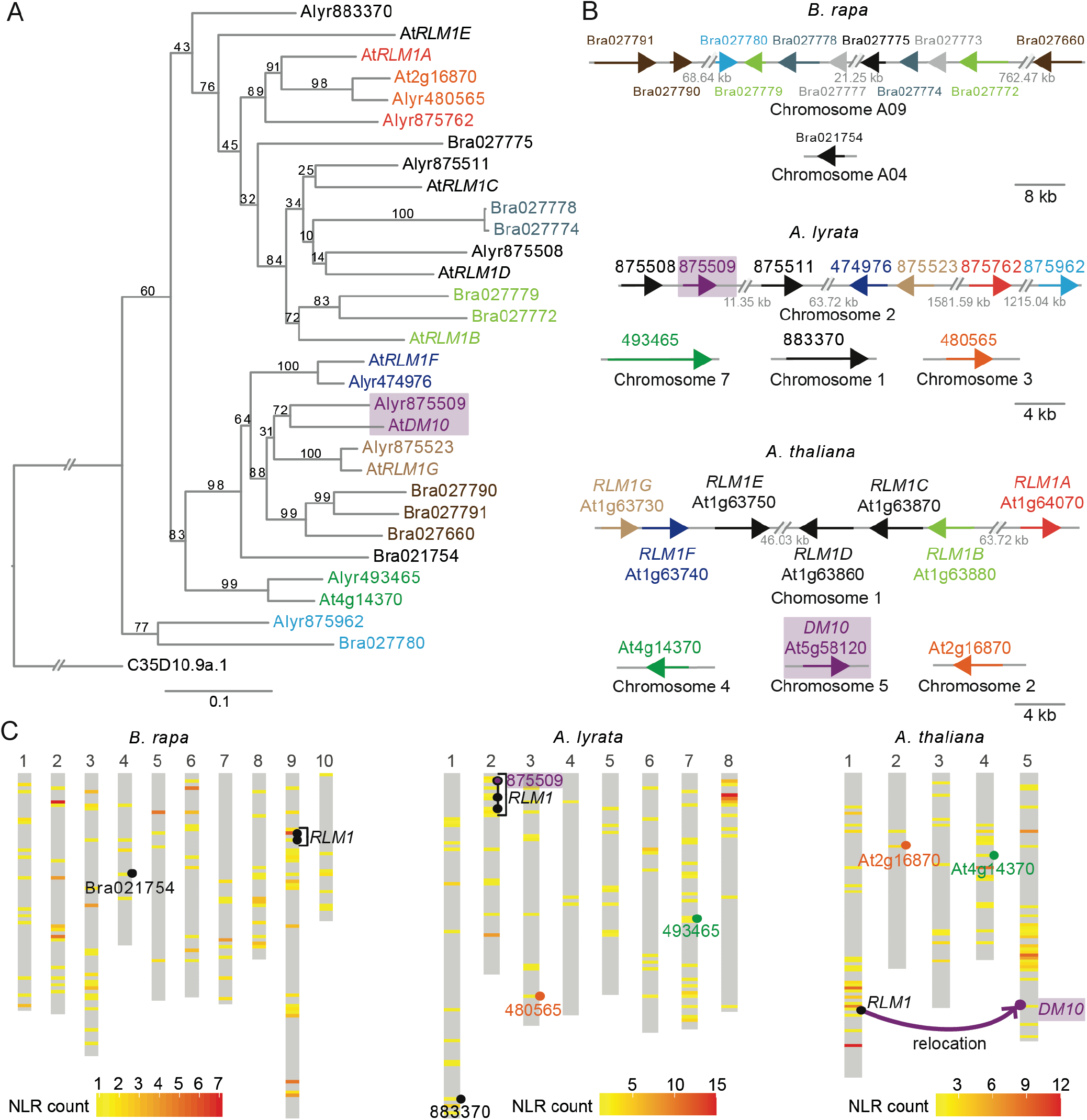
*RLM1* locus in *B. rapa*, *A. lyrata* and *A. thaliana* reference genomes. **A.** ML tree of the NBS domain (CDS) of *RLM1* cluster members from *B. rapa*, *A. lyrata* and *A. thaliana*, with the NBS domain of *C. elegans* CED-4 as outgroup (C35D10.9a.1). 1,000 bootstrap replicates were performed, values are shown on each branch. Branch lengths in nucleotide substitutions are indicated. The same color was chosen for genes in neighboring branches with bootstrapping values above 70. Diagonal lines indicate a gap in the tree branches. **B.** *RLM1* cluster members in *B. rapa*, *A. lyrata* and *A. thaliana*. Color-coding the same as in A, genes in grey are truncated, arrows represent size of NLR loci. Diagonal lines indicate a positional gap along the chromosome, the length of the gap is indicated. **C.** Heatmaps of NLR densities across the three genomes. Window sizes were calculated by dividing the length of the longest chromosome by 100. *RLM1* cluster and closely related singletons indicated. *DM10* in *A. thaliana* and its homolog in *A. lyrata*, 875509 are highlighted in magenta.

## Discussion

Over ten causal genes for hybrid necrosis have been identified in *A. thaliana* and other plants (Krüger et al. 2002; Bomblies et al. 2007; Alcázar et al. 2009; Jeuken et al. 2009; Yamamoto et al. 2010; Chae et al. 2014; Chen et al. 2014; Todesco et al. 2014; Sicard et al. 2015; Deng et al. 2019). In many instances, at least one of the two causal genes is an NLR, which is also the case for the Cdm-0 x TueScha-9 incompatibility. What makes this case particularly interesting is the extreme severity of hybrid necrosis, the transcriptional hyper-induction of NLR genes, and the causality of a truncated singleton NLR, *DM10*, which was recently relocated from a larger NLR cluster.

As with normal immune responses and autoimmune syndromes, the expression of hybrid necrosis is typically temperature-dependent, and hybrid necrosis in *A. thaliana* can usually be completely suppressed when grown above 23°C (Bomblies et al. 2007; Alcázar et al. 2009; Chae et al. 2014; Todesco et al. 2014; Świadek et al. 2017). In contrast, Cdm-0 x TueScha-9 F_1_ seedlings die even at 28°C (Chae et al. 2014). In other necrotic hybrids, non-causal NLRs have been reported to be differentially expressed between hybrids and their parents (Bomblies et al. 2007), but not anywhere near the massive NLR induction seen in Cdm-0 x TueScha-9. In the F_1_ progeny, 128 of 150 expressed NLRs are differentially expressed in at least one genotype comparison, with almost all being overexpressed. Simultaneous upregulation of several NLR genes has been observed after exposure to biotic (Zipfel et al. 2004; Tan et al. 2007; Ribot et al. 2008; Mohr et al. 2010; Yu et al. 2013; Sohn et al. 2014; Chen et al. 2015; Mine et al. 2018) and abiotic stresses (MacQueen and Bergelson 2016), but also not to the extent seen in Cdm-0 x TueScha-9 hybrids. Given that elevated NLR expression levels can trigger cell death (Stokes et al. 2002; Mackey et al. 2003; Palma et al. 2010; Lai and Eulgem 2018), we expect that NLR hyper-induction is a significant contributor to the strongly necrotic phenotype of Cdm-0 x TueScha-9 F_1_ hybrids.

NLR transcript levels are tightly controlled through a variety of regulatory mechanisms (Lai and Eulgem 2018), and large-scale upregulation of NLRs could possibly require multiple pathways. We found WRKY transcription factors to be overexpressed in the hybrid; these proteins bind to W box motifs enriched in the promoters of multiple members of the plant immune system, including NLRs, and can induce widespread NLR expression, enhancing basal immunity (Eulgem and Somssich 2007; Pandey and Somssich 2009; Mohr et al. 2010). Two other mechanisms known to affect a broad set of NLRs is the miRNA-dependent phasiRNA production (Zhai et al. 2011; Li et al. 2012; Shivaprasad et al. 2012; Xia et al. 2015) as well as nonsense-mediated decay (NMD) (Gloggnitzer et al. 2014), both of which help to dampen NLR gene expression in the absence of pathogen threats. Repression is attenuated after an incoming pathogen is detected by the plant, enabling global NLR levels to increase (Lai and Eulgem 2018). While we have no direct evidence for transcription factors, small RNAs or NMD as contributors to aberrant NLR expression in the Cdm-0 x TueScha-9 hybrid, this exceptional hybrid necrosis case may present a good tool for comparing NLR regulation under pathogen attack with strong autoimmunity.

Allelic variation in singleton NLRs can manifest itself as presence/absence polymorphisms, intragenic recombination events or single nucleotide changes (Caicedo et al. 1999; Stahl et al. 1999; Mauricio et al. 2003; Allen et al. 2004). Truncations may result, for example, from point mutations (Cao et al. 2011; Guo et al. 2011), alternative splicing or transcriptional termination (Costanzo and Jia 2009; Kim et al. 2009; Tsuchiya and Eulgem 2013). We found 17% of DM10 proteins encoded in a global set of *A. thaliana* accessions to be truncated in either their TIR, NBS or LRR domain. Similar to several full-length variants, the alleles for all three truncated proteins have intermediate frequencies and are relatively wide-spread, suggesting they are actively maintained in the global population by balancing selection. The most common of the three truncation alleles is the *DM10* risk version, which has evidence for its LRR domain being under diversifying selection. While the TIR domain alone can induce cell death (Swiderski et al. 2009; Bernoux et al. 2011), a complete NBS domain is essential in many instances (Dinesh-Kumar et al. 2000; Dodds et al. 2001; Bendahmane et al. 2002; Tameling et al. 2002; Williams et al. 2011; Steinbrenner et al. 2015; Sueldo et al. 2015; Wang et al. 2015; Bernoux et al. 2016). Thus, it is unclear whether the shorter DM10^Cdm-0^ or DM10^Sha^ proteins are functional, even though their prevalence and geographical distribution suggest so.

Because NLR allelic diversity is often not easily captured by short-read based resequencing (Van de Weyer et al. 2019), we still do not have a good grasp on whether NLR alleles in general, and specifically beneficial alleles, spread through the population more quickly than other adaptive alleles. The Iberian peninsula is a center of *A. thaliana* genetic diversity, with strong geographical structure across a north-south latitudinal gradient (Picó et al. 2008; Brennan et al. 2014). We observed a lack of co-occurrence between *DM11* risk alleles, restricted to southwestern Spain, and *DM10* risk alleles, restricted to the northern half of Spain (1001 Genomes Consortium 2016). Absence of co-occurrence between risk alleles may partly be the result of population structure: two geographical barriers potentially reducing gene flow, the Tagus river and the Central System mountains, divide populations carrying either *DM10* or *DM11* risk alleles. In any case, more definitive proof of the mutual exclusion of *DM10* and *DM11* risk alleles will require more extensive sampling of natural populations across the Iberian peninsula. Co-occurrence of hybrid incompatibility alleles in a single population has been observed before, where different alleles are maintained at intermediate frequencies, but in this case, the hybrids show a milder necrosis phenotype in the lab than Cdm-0 x TueScha-9, and no obvious phenotype in the wild (Todesco et al. 2014). The extreme necrotic phenotype caused by the *DM10*-*DM11* interaction, which appears to be largely independent of growth conditions, makes it unlikely that the hybrid phenotype would be suppressed in the wild. In addition, since outcrossing rates of *A. thaliana* in the wild can be substantial (Bomblies et al. 2010), it is conceivable that in some areas these rates are high enough for lethal hybrids to exact a noticeable fitness cost on risk allele carriers.

An interchromosomal relocalization event of the *RLM1* cluster gave rise to *DM10* after *A. thaliana* speciation. Which evolutionary forces might have caused *DM10* to be relocated to a separate chromosome, if any? NLR genes in clusters are likely to be more mutable than singletons because of illegitimate recombination (Michelmore and Meyers 1998; Baumgarten et al. 2003; Meyers et al. 2005; Wong and Wolfe 2005; Wicker et al. 2007). If *DM10* underwent beneficial neofunctionalization after duplication, its relocation away from the cluster might have stabilized the locus. Another possibility could be conflicts among gene cluster members. Cluster members are sometimes transcriptionally co-regulated (Yi and Richards 2007; Deng et al. 2017), so translocation away from the cluster would allow for evolution of new expression patterns for *DM10*. More generally, genomic relocation would enable *DM10* to be subjected to different selection regimes than its cluster homologs. Either way, the fact that the genomic region surrounding *DM10* – different from some other *RLM1* cluster members – is a recombination cold spot (Choi et al. 2016) is consistent with our finding of high LD around the *DM10* locus, especially in accessions carrying the *DM10* risk allele. Together with our phylogenetic results and Tajima’s *D* measurements, this would seem to support the idea of stable *DM10* haplotypes being particularly advantageous.

In conclusion, we have presented a severe case of hybrid necrosis in *A. thaliana*, where the hybrid showed global NLR hyper-induction triggered by the interaction of *DM10*, a relocated singleton NLR, and *DM11*, an unliked locus in chromosome 1. It is noteworthy that the *DM10* risk alleles lack a substantial portion of the coding region, including sequences for three of the eight LRRs found in the full-length variants. NLRs lacking the NBS or LRR domain are not only known to retain the ability to cause cell death, but there are cases where truncated NLRs are bona fide resistance genes (Nishimura et al. 2017; Roth et al. 2017; Marchal et al. 2018). Conversely, other proteins, including at least one full-length NLR, can induce cell death through activation of naturally occurring truncated NLRs (Zhao et al. 2015; Zhang et al. 2017). In the case of *DM10*, we do not know whether only the full-length variants or the truncated variants, or both, confer resistance to unknown pathogens. To answer this question, a new program is needed that systematically assigns function to all NLR alleles and genes in a species. Regardless of whether the truncated *DM10* alleles have such activity, already with our current knowledge, the *DM10*/*DM11* case provides a good tool to investigate the consequences of simultaneous activation of a large fraction of NLRs. In the future, by identifying the role of different alleles of *DM10* and its homologs in the *RLM1* cluster in responses to natural pathogens, one could test whether chromosomal relocation affects how evolution is acting on this group of highly related NLR genes.

## Materials and Methods

### Constructs are listed in Table S15 and primers in Table S16

Sequencing data can be found at the European Nucleotide Archive (ENA) under accession number PRJEB38267 (RNA-seq experiment) and ### (Cdm-0 assembly) and in the GenBank under accession number ### (*DM10* region).

### Plant material

Stock numbers of accessions used are listed in Supplementary Material. All plants were stratified in the dark at 4°C for 4-6 days prior to planting on soil. Late flowering accessions were vernalized six weeks under short day conditions (8 h light) at 4°C. All plants were grown in long days (16 h of light) at 16°C or 23°C at 65% relative humidity under 110 to 140 μmol m^−2^ s^−1^ light provided by Philips GreenPower TLED modules (Philips Lighting GmbH, Hamburg, Germany).

### RNA sequencing

Six biological replicates of 10 day-old shoots of Cdm-0 x TueScha-9 hybrids and their parental accessions were collected. RNA was extracted as described in (Yaffe et al. 2012). The NEBNext magnetic isolation module (New England Biolabs), was used for mRNA enrichment. Sequencing libraries were prepared using NEBNext Ultra II directional RNA library kit and paired-end sequenced (150bp) in an Illumina HiSeq3000 (Illumina Inc., San Diego, USA) instrument. Reads were mapped against the *A. thaliana* reference TAIR10 using bowtie2 (v.2.2.6) (Langmead and Salzberg 2012). Default parameters were chosen unless mentioned otherwise. Transcript abundance was calculated with RSEM (v.1.2.31) (Li and Dewey 2011). In silico hybrids were generated to enable mid-parent value calculations: parental read files were normalized according to sequencing depth and were subsampled by randomly drawing 50 % of the reads with seqtk (v.2.0-r82-dirty; https://github.com/lh3/seqtk). Differential gene expression analyses were performed using DESeq2 (v.1.18.1) (Love et al. 2014). Genes with less than ten counts over all 18 samples were removed from downstream analyses. Significant changes in gene expression between two genotypes were determined by filtering for genes with a |log_2_FoldChange| >1 and padj value < 0.01. One read was added to all normalized read counts in Fig 1G and S2E to avoid plotting-INF values in non-expressed genes (log_10_(0+1)=0). Non-additive gene expression between Cdm-0 x TueScha-9 F_1_ hybrids in silico hybrids was analyzed by computing principal components based on the normalized read counts of the top 500 most variable genes across all 18 samples. Plots were generated using the R package ggplot2 (v.3.2.0) (Wickham 2009) and heatmaps were plotted using pheatmap (v.1.0.8) (Kolde 2012). Gene Ontology (GO) analyses were performed using AgriGO (Tian et al. 2017) using the SEA method. The GO results were visualized with REVIGO treemap (Supek et al. 2011), for clearer visualization only the top 13 GO categories with the lowest −log_10_(*p* value) were plotted, the complete list of GO terms is found in Table S2.

### Genotyping-by-sequencing and QTL mapping

F_1_ progeny from bi-directional crosses of F_1_ (TueScha-9/Col-0) x (Cdm-0/Col-0) was used as a mapping population. The seedlings showing the hybrid necrosis phenotype vs. those that did not, were genotyped individually in a 1:1 ratio. Plants were 10 days old when collected. Genomic DNA was extracted with CTAB (cetyl trimethyl ammonium bromide) buffer (Doyle and Doyle 1987) and then purified through chloroform extraction and isopropanol precipitation (Ashktorab and Cohen 1992). Genotyping-by-Sequencing (GBS) using RAD-seq was used to genotype individuals in the mapping populations with KpnI tags (Rowan et al. 2017). Briefly, libraries were single-end sequenced on a HiSeq 3000 instrument with 150 bp reads. Reads were processed with Stacks (v1.35) (Catchen et al. 2013) and mapped to TAIR10 with bwa-mem (v0.7.15) (Li 2013), variant calling was performed with GATK (v3.5) (McKenna et al. 2010). QTL was performed using R/qtl (Broman et al. 2003) with the information from 348 F_2_ individuals from 4 independent lines of this segregating population and 6179 markers.

### *De novo* genome assembly and annotation

The Cdm-0 accession (ID 9943; CS76410) was grown as described above. To reduce starch accumulation, 3-week-old plants were put into darkness for 30 h before harvesting. Sixteen grams of flash frozen leaf tissue were ground in liquid nitrogen and nuclei isolation was performed according to (Workman et al. 2018) with the following modifications for *A. thaliana*: eight independent reactions of two grams each were carried out, and the filtered cellular homogenate was centrifuged at 7,000 x g. High-molecular-weight DNA was recovered with the Nanobind Plant Nuclei Kit (Circulomics; SKU NB-900-801-01), and needle-sheared 1x (ThermoFisher UK Ltd HCA-413-030Y GC Syringe Replacement Parts 26g, 51mm). A 35-kb template library was prepared with the SMRTbell® Express Template Preparation Kit 2.0, and size-selected with the BluePippin system according to the manufacturer’s instructions (P/N 101-693-800-01, Pacific Biosciences, California, USA). The final library was sequenced on a Pacific Biosciences Sequel instrument with Binding Kit 3.0. PacBio long-reads were assembled with Canu (v1.71) (Koren et al. 2017). The resulting contigs were first polished using the long-reads with the Arrow algorithm (v2.3.2; https://github.com/PacificBiosciences/GenomicConsensus), followed by a second polishing step with PCR-free short-reads using the Pilon algorithm (v1.22) (Walker et al. 2014). Lastly, the resulting contigs were scaffolded based on TAIR10 assembly by REVEAL (v0.2.1) (Linthorst et al. 2015). The previously generated Cdm-0 transcriptome sequencing data were mapped against the scaffolded genome assembly using HISAT (v.2.0.5) (Kim et al. 2015)). Subsequently, the mapping results were used as extrinsic RNA sequencing evidence when annotating the genome using AUGUSTUS (v3.2.3) (Stanke et al. 2006).

### Manual NLR annotation of the DM11 mapping interval

The 20-25 Mb region of chromosome 1 was extracted from the Cdm-0 assembly. The assembly was used as a query against a subject FASTA file containing 167 NLR genes from the Col-0 reference accession using blastn (Altschul et al. 1990). Hits were binned in 20 kb intervals and the percentage identity between the queries and the subject was visualized across all bins. NLRs between At1g56510 to At1g64070 in Col-0 found in this interval were manually annotated based on the percentage identity plotted and on AUGUSTUS gene predictions (v2.5.5) (Stanke et al. 2006).

### GWAS

Cdm-0-dependent hybrid necrosis in the F_1_ progeny from crosses with 80 accessions (Chae et al. 2014) was scored as 0 or 1. The binary trait with accession information was submitted to the easyGWAS platform (Grimm et al. 2016) using the FaSTLMM algorithm. A −log_10_(*p* value) was calculated for every SNP along the five *A. thaliana* chromosomes.

### Constructs and transgenic lines

Genomic fragments were PCR amplified, cloned into pGEM®-T Easy (Promega, Madison, WI, USA) and then transferred to the binary vector pMLBart, pCambia1300 or pFK210. Constructs were introduced into plants using *Agrobacterium*-mediated transformation (Weigel and Glazebrook 2002). T_1_ transformants were selected on BASTA (pMLBart and pFK210) and crossed to incompatible accessions. Ethanol-inducible constructs were PCR amplified, cloned into pGEM®-T Easy, as part of a separate experiment, 2xHA tags were added via PCR and the whole fragment, which was then transferred to the pCR8® entry vector (ThermoFisher Scientific). Next, the genomic fragment was moved to the destination vector pZZ006 (Caddick et al. 1998) through the Gateway® LR reaction (ThermoFisher Scientific). Quality control for all constructs was done by Sanger sequencing. For transient expression in *N. benthamiana, A. tumefaciens* were grown to an OD_600_ of 1.2-1.8, incubated in induction medium (10 mM MES (pH 5.6), 10 mM MgCl2, and 150 μM acetosyringone). The cell suspensions were normalized to an OD_600_ of 0.8 and co-infiltrations suspensions were mixed 1:1. Suspensions were then infiltrated into the abaxial side of *N. benthamiana* leaves. In the case of EtOH inducible constructs, infiltrated *N. benthamiana* were induced at 18 h post-infiltration (hpi) by irrigation with 1% ethanol and kept within a transparent plastic dome for another 18 h. *DM10 N. benthamiana* constructs were co-expressed with a *35S::GFP* construct as part of a larger experiment to test for candidate *DM11* loci.

### Population genetic analyses

Amino acid sequence conservation scores were calculated with ConSurf (Armon et al. 2001; Ashkenazy et al. 2016). SNPs occurring in repetitive regions and only present in one of the 73 extracted *DM10* alleles were considered sequencing errors and were manually curated. Protein domains were predicted using InterProScan (Jones et al. 2014). LRR domains were predicted with LRRsearch and the score threshold was set at 7 (Bej et al. 2014). NLR motifs were defined based on previous studies (Meyers et al. 2003; Shao et al. 2016). Nonsynonymous to synonymous substitution rates were calculated using KaKs_Calculator (v2.0) (Wang et al. 2010) with the NG method (Nei and Gojobori 1986); a window length of 60 bp and a step size of 6 bp were chosen. Genomic regions of interest were subsetted from a 1135 genomes VCF file (1001 Genomes Consortium 2016) using VCFtools (v0.1.14) (Danecek et al. 2011). The resulting VCF file was filtered by MAF=0.01 and a maximum percent of missing data per SNP of 30%. Sequences were converted to FASTA, aligned with MUSCLE (v3.8.31) (Edgar 2004) and then visualized with Aliview (v1.18.1) (Larsson 2014). Neighbor-Joining trees were calculated with Fastphylo (v1.0.1) (Khan et al. 2013) and visualized with iTol (Letunic and Bork 2007). Maximum-likelihood trees were calculated with RaxML (v.0.6.0) using the GTR+G4 model (Stamatakis 2014). Linkage disequilibrium (r^2^), was calculated with PLINK (v.1.90) (Purcell et al. 2007). Principal component analyses were calculated with smartPCA (Patterson et al. 2006). Tajima’s *D, F_ST_*, and nucleotide diversity (**π**) were also calculated with VCFtools. Maps were created with the R-packages maps (v3.3) and ggmap (v3.0) (Kahle and Wickham 2013). Admixture groups were assigned to each accession in accordance with the 1001 Genomes project (1001 Genomes Consortium 2016); since TueScha-9 had not been part of that study, admixture group assignments for it were estimated based on the genetic make-up of neighboring accessions. *RLM1* homologs in *A. lyrata* and *B. rapa* were identified using the Ensembl Plants portal (Bolser et al. 2016). Sequences from the genome assemblies TAIR10 (*A. thaliana*), *B. rapa* (v1.5) and *A. lyrata* (v1.0) were used for phylogenetic analyses.

## Author contributions

**Conceptualization:** ACB, DW, EC.

**Formal analysis:** ACB, MC, RL, FR, HA, EC.

**Funding acquisition:** DW, EC.

**Investigation:** ACB, JW, WYC, EC.

**Methodology:** ACB, EC.

**Project administration:** DW.

**Supervision:** DW.

**Writing – original draft:** ACB.

**Writing – review & editing:** ACB, DW, EC.

## Competing interests

The authors have declared that no competing interests exist.

## Funding

This work was supported by the ERC Advanced Grant IMMUNEMESIS (340602), the Deutsche Forschungsgemeinschaft through the Collaborative Research Center (CRC1101), the Max Planck Society (to D.W.) and the Academic Research Fund (MOE2019-T2-1-134) from the Ministry of Education, Singapore, Intramural Research Fund (R-154-000-B33-114) from the National University of Singapore (to E.C.). The funders had no role in study design, data collection and analysis, decision to publish, or preparation of the manuscript.

## Acknowledgements

We thank Sang-Tae Kim and members of the Weigel lab for critical reading of the manuscript, Christa Lanz for preparing the Cdm-0 PacBio library, Lei Li for performing the *RLM1 N. benthamiana* transient expressions, Rebecca Schwab for generating a few crosses as well as Gautam Shirsekar and Sergio Latorre for discussion. We also thank the 1001G+ team for providing access to preliminary whole genome *A. thaliana* assemblies.

## Notes

### Competing Interest Statement

The authors have declared no competing interest.

