## Supplemental Figures for "A singleton NLR of recent origin causes hybrid necrosis in *Arabidopsis thaliana*"

**Fig S1. RNA-seq analysis of Cdm-0 x TueScha-9 hybrid plants.** Related to Fig 1.

**Fig S2. Identification of *DM10* and *DM11*.** Related to Fig 2.

**Fig S3. *De novo* Cdm-0 genome assembly.** Related to Fig 2.

**Fig S4. Pairwise genetic distances for three *DM10* candidate genes across 80 accessions.** Related to Fig 3.

**Fig S5. *DM10* natural variation.** Related to Fig 5.

### Supplemental Methods

### Supplemental References

### Supplemental Figures

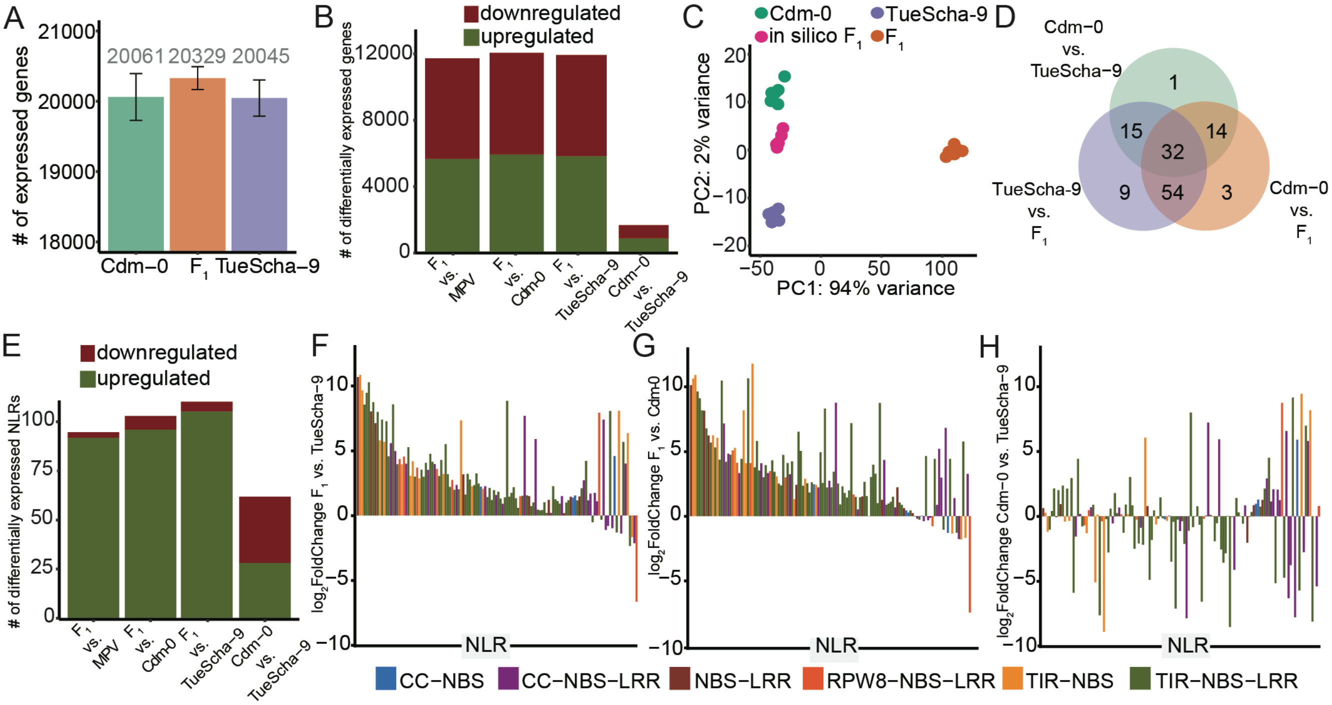


**Fig S1. RNA-seq analysis of Cdm-0 x TueScha-9 hybrid plants. A.** Total number of expressed genes in both the $F_1$ hybrid and parents. **B.** Significantly (|log$_2$FoldChange| >1, padj value < 0.01) up- and downregulated genes across different genotype comparisons. **C.** PCA of gene expression variance separating the $F_1$ hybrids, parents and in silico hybrids. **D.** Intersection of differentially expressed NLRs between the $F_1$ hybrid and parents. **E**. Significantly up- and downregulated NLR genes across different genotype comparisons. **F-H.** NLR expression changes between the $F_1$ hybrid and TueScha-9 (F), $F_1$ hybrid and Cdm-0 (G), Cdm-0 and TueScha-9 (H). The NLR gene order follows Fig 1G.

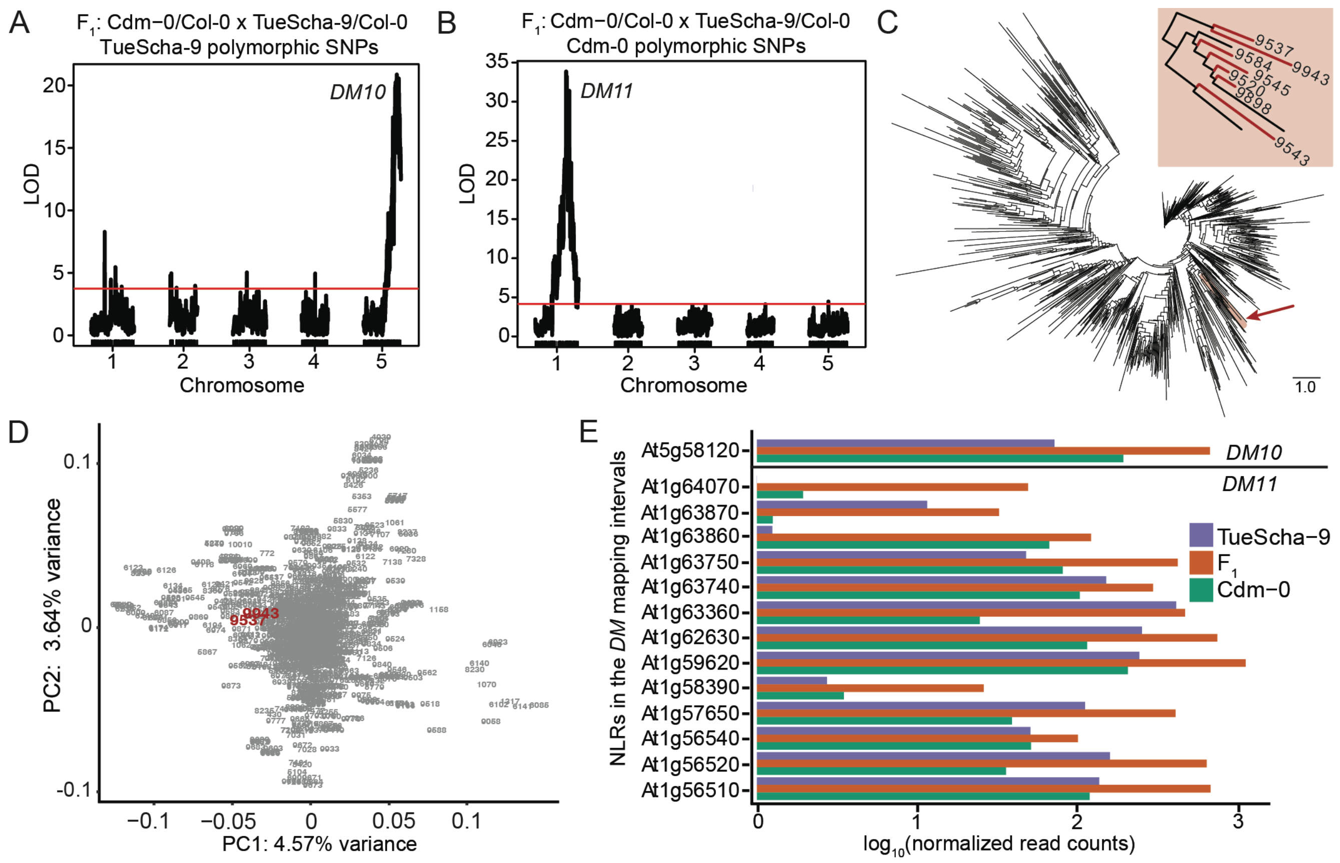


**Fig S2. Identification of *DM10* and *DM11*. A.** Polymorphic SNPs from TueScha-9, *DM10* on chromosome 5 (22.35-25.27 Mb). **B.** Polymorphic SNPs from Cdm-0, *DM11* on chromosome 1 (21.55-22.18 Mb). Horizontal lines indicate 0.05 significance threshold established with 1,000 permutations. **C.** Example of an NJ tree from one of the *DM11* candidate loci: At1g59780. Region where Cdm-0 (9943) and IP-Cum-1 (9537) are found is highlighted in red. Inset shows a close-up of Cdm-0-like accessions, accessions crossed to TueScha-9 are marked in red. **D.** Example of a PCA plot using VCF information for the entire *DM11* mapping interval from the 1001 Genomes Project (1001 Genomes Consortium 2016). Accession IDs from the 1001 Genomes Project in grey, with Cdm-0 (9943) and IP-Cum-1 (9537) in red. **E.** Normalized RNA-seq read counts for the hybrid and parents. Shown are the only NLR in the *DM10* mapping interval, At5g58120, as well as NLRs found between At1g56510 and At1g64070 on chromosome 1, which includes the *DM11* interval. Missing bars mean the gene was not expressed in that genotype.

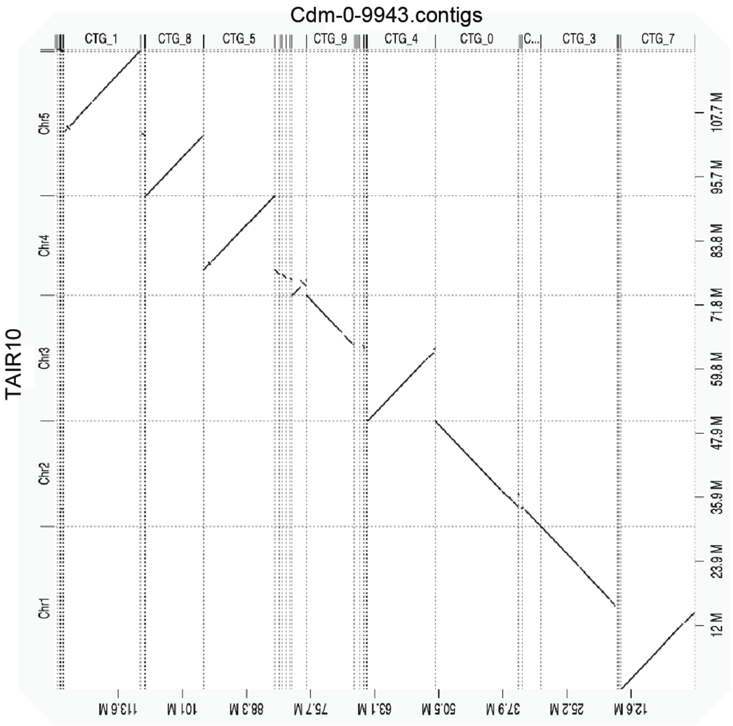


**Fig S3. *De novo* Cdm-0 genome assembly.** Dot plot based on minimap2 (Li 2018) alignment between the Cdm-0 contigs and the reference genome (TAIR10) using D-GENIES (Cabanettes and Klopp 2018).

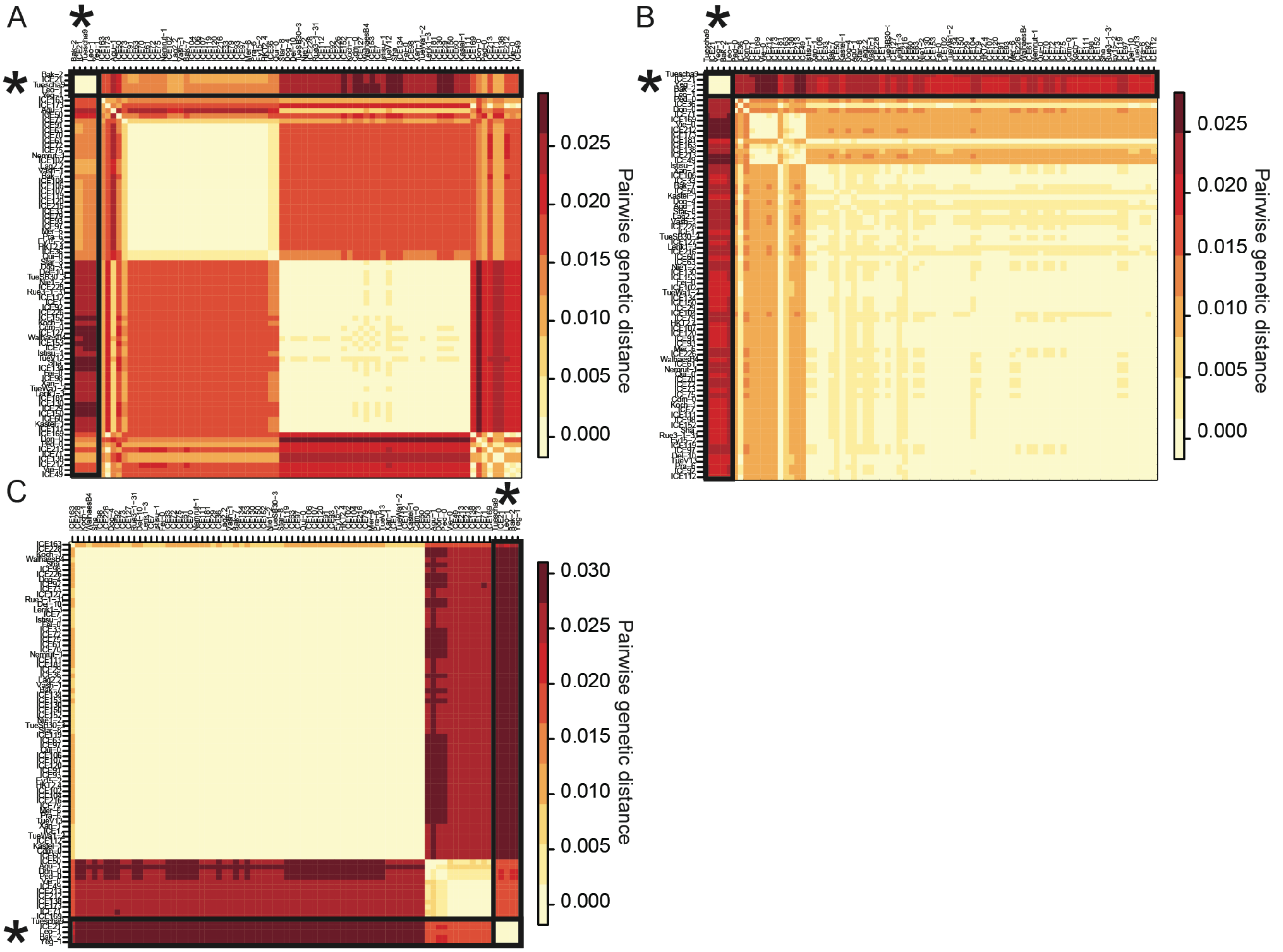


**Fig S4. Pairwise genetic distances for *DM10* three candidate genes across 80 accessions. A-C.** Heatmaps of pairwise genetic distances among 80 alleles of At5g58120 (A), *ROS3* (B) and *PHOT2* (C). Distances are the fraction of nucleotide sites at which two sequences are different. Asterisk highlights the group of five risk accessions causing hybrid necrosis when crossed to Cdm-0. These five accessions are genetically very similar in all three genes.

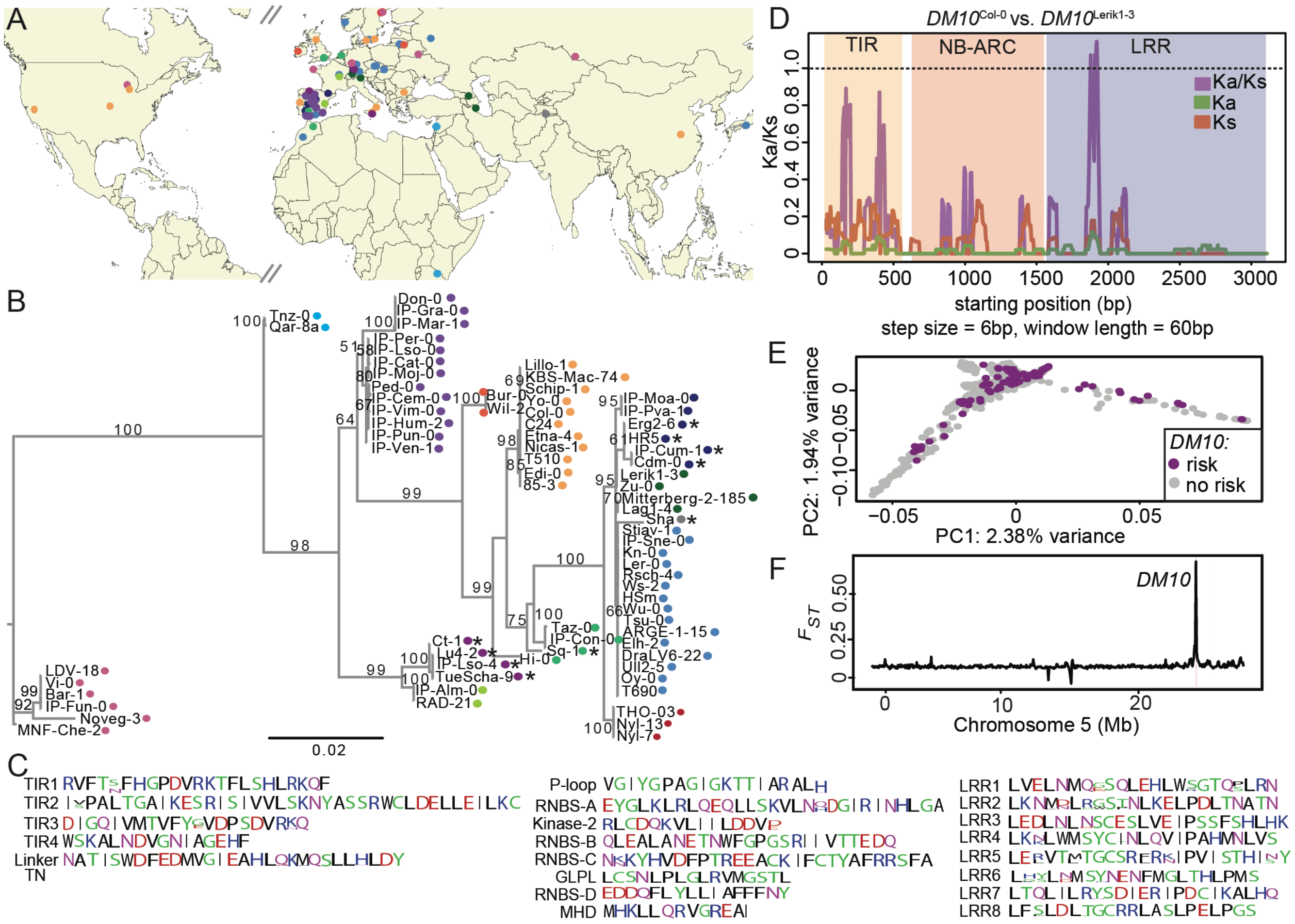


**Fig S5. *DM10* natural variation. A.** Geographic locations of 73 accessions carrying different *DM10* alleles. Each color indicates a similar *DM10* allele. **B.** ML tree of 73 CDS *DM10* sequences. 1,000 bootstrap replicates were performed, bootstrapping values are indicated on each branch, values above 50 are shown. Branch lengths in nucleotide substitutions are indicated. Asterisks indicate truncated DM10 proteins, colors as in A. **C.** DM10 motif consensus across 73 accessions. **D.** Ka/Ks ratio between $DM10^{Col-0}$ and $DM10^{Lerik1-3}$ **E.** Whole-genome PCA of 1001 Genomes accessions. **F.** $F_{ST}$ between *DM10* risk and non-risk accessions across chromosome 5, only one peak is found in the region where *DM10* is located.

### Supplemental Methods

Pairwise genetic distances were calculated using the dist.DNA function in the ape R-package (v5.2) (Paradis and Schliep 2019). The identified NLR motif consensus across 73 DM10 proteins was visualized using WebLogo (Crooks et al. 2004).
